# Targeted proteomics addresses selectivity and complexity of protein degradation by autophagy

**DOI:** 10.1101/2024.03.27.586977

**Authors:** Alexandre Leytens, Rocío Benítez-Fernández, Carlos Jiménez-García, Carole Roubaty, Michael Stumpe, Patricia Boya, Jörn Dengjel

## Abstract

Autophagy is a constitutively active catabolic lysosomal degradation pathway, often found dysregulated in human diseases. It is often considered to act in a cytoprotective manner and is commonly upregulated in cells undergoing stress. Its initiation is regulated at the protein level and does not require *de novo* protein synthesis. Historically, autophagy has been regarded as non-selective; however, it is now clear that different stimuli can lead to the selective degradation of cellular components via selective autophagy receptors (SARs). Due to its selective nature and the existence of multiple degradation pathways potentially acting in concert, monitoring of autophagy flux, *i.e.* selective autophagy-dependent protein degradation, should address this complexity. Here, we introduce a targeted proteomics approach monitoring abundance changes of 37 autophagy-relevant proteins covering process-relevant proteins such as the initiation complex and the ATG8 lipidation machinery, as well as most known SARs. We show that proteins involved in autophagosome biogenesis are upregulated and spared from degradation under autophagy inducing conditions in contrast to SARs. Classical bulk stimuli such as nutrient starvation mainly induce degradation of ubiquitin-dependent soluble SARs and not of ubiquitin-independent, membrane-bound SARs. In contrast, treatment with the iron chelator deferiprone leads to the degradation of ubiquitin-dependent and - independent SARs linked to mitophagy and reticulophagy/ER-phagy. Our approach is automatable and supports large-scale screening assays paving the way to (pre)clinical applications and monitoring of specific autophagy fluxes.

## Introduction

Eukaryotic cells maintain homeostasis and remove damaged or superfluous cellular components through a conserved degradation pathway called macroautophagy (hereafter referred to as autophagy). This highly conserved catabolic pathway occurs at basal levels but is enhanced by a variety of stress signals, such as nutrient starvation or organelle damage (1). The process starts with the *de-novo* formation of a double-membrane organelle called autophagosome from membranes being recruited from various sources, the endoplasmic reticulum (ER) likely being principal donor (2). This growing cup-shaped membrane engulfs parts of the cytoplasm and closes to form a double membrane vesicle. The autophagosome can fuse with other vesicles from endocytic pathways forming an amphisome and finishes by fusing with the lysosome/vacuole. Lysosomal fusion exposes autophagosomal cargo including the inner membrane to acidic hydrolases and enables the degradation of its content. The generated building blocks are recycled and transported back to the cytosol by lysosomal permeases to generate energy or fuel anabolic pathways.

While basal autophagy is often considered a “bulk”, *i.e.* a non-selective process recycling cellular components in a random manner, autophagy can also be selective. Well studied cases of selective autophagy include the targeted removal of damaged organelles such as the mitochondrion or ER (termed mitophagy and reticulophagy/ER-phagy, respectively) as well as of protein aggregates (aggrephagy). This particular selectivity is enabled by a set of proteins called selective autophagy receptors (SARs) (3), bridging the autophagy cargo to proteins of the ATG8 family, which are lipidated proteins coating the autophagosomal membrane and functioning as docking sites (4).

Numerous methods and protocols have been described for the study of protein turnover by autophagy, several of them relying on immunodetection or fluorescent microscopy of single proteins, which are often ectopically expressed as tagged variants facilitating downstream analyses (5). While these methods are well established, they often require a significant protein amount or protein tagging, making them unsuitable for some sample types, *e.g.* primary cells being difficult to transfect, and difficult to adapt for large-scale screening approaches. Additionally, most of these methods infer the activity of the whole pathway based on the quantification of single markers. Whereas this might be relevant for the analysis of basal, *i.e.* non-selective, autophagy flux, such approaches likely fail to capture the complexity of stress-induced autophagy in which different cargoes might get degraded at different rates and through different sub-pathways (6). To address the complexity of autophagy regulation and activity in an unbiased manner, approaches monitoring expression of multiple genes by RNA-seq have been developed that are adaptable for higher sample throughput and might also be used in prognostic/diagnostic settings (7). However, to infer autophagy activity based on changes in gene transcription biological samples have to be treated for several hours, which is in contrast to the rapid cellular response to stress conditions. Autophagy signaling is activated within minutes (8), initial autophagosome biogenesis does not require *de novo* protein synthesis (9), and in mammalian cells autophagosomes and autophagy-dependent protein degradation can be detected as early as 30 min post stimulus (10). Additionally, while transcriptomic approaches are well suited to investigate future adaptations of the cells to a given stimulus, they cannot answer questions about protein degradation itself. To address this gap in analytical approaches, we developed an assay relying on detecting changes in protein abundances, which is compatible with high sample throughput, and which monitors abundance changes of multiple endogenous proteins reflecting the complexity and multitude of autophagy subtypes.

We turned to targeted proteomics, which are mass spectrometry (MS)-based analyses enabling the accurate quantification of a predefined number of given peptides potentially encompassing numerous target proteins (11), supporting absolute protein quantification (12), and being highly sensitive with limits of detection in the Attomole range. Mass spectrometers operated in targeted mode focus on a pre-established set of mass-to-charge ratios corresponding to peptides-of-interest. By ignoring other peptides eluting from the coupled liquid chromatography (LC) column, targeted proteomics greatly enhances the sensitivity for its targets. Combining this approach with labelled spiked-in synthetic peptides further enables confident identification of the targets, robust normalization across samples, and absolute quantification, depending on peptide purity (13). Here, we describe a targeted proteomics approach based on parallel reaction monitoring (PRM) (14), allowing to accurately quantify tryptic peptides derived from human autophagy-relevant proteins in single MS measurements. In combination with a simple in-solution digestion protocol that can be performed in a (semi)automated manner the assay supports screening approaches. We monitor proteins involved in the autophagic responses to amino acid and glucose starvation, as well as to deferiprone (DFP) treatment, a potent inducer of mitophagy (15), covering autophagy initiation, autophagosome biogenesis and turnover, as well as SARs enabling an accurate assessment of the activity and selectivity of the entire process.

## Results

### Target selection and peptide characterization

To cover bulk and selective autophagy subtypes by targeted proteomics, we screened the respective literature for relevant proteins (1, 4, 7, 16). In addition, we included selected, autophagy-relevant proteins from ongoing research projects in our groups (17), which led to a list of 41 proteins-of-interest (POIs) (**supplemental Table S1**). MS-compatible peptides of POIs for targeted proteomics are commonly generated by proteolytic digestion using the endoprotease trypsin. Not all tryptic peptides from a given source protein are suited for targeted MS as some sequences might not fit the requirements of LC (highly hydrophilic or hydrophobic peptides), chemical stability (*e.g.* post-lysis methionine oxidation), or distribution of trypsin cleavage sites (lysine or arginine positions in the sequence prevent digestion into suitable peptides yielding either too large or too small peptides). For these reasons, only peptide sequences between 7 and 25 amino acids in lengths were considered. Due to the sensitivity to oxidations, methionine-containing sequences were excluded. Unfortunately, these restrictions led to the exclusion of some important protein candidates. As an example, human ATG8 proteins are very short proteins that only generate a limited number of tryptic peptides. Among this small set, some peptides had to be removed as their sequences did not match our quality criteria. As a result, only three peptides from GABARAPL1 and GABARAPL2 were further considered and no LC3-derived peptides (MAP1LC3A, B, B2, C) made it to the final list. As a next step, shortlisted peptides were screened on their detection in previous MS experiments in publicly available MS datasets from PeptideAtlas as well as by our group (18). This led to a selection of 114 peptides sequences from the 41 POIs. These were synthesized following the AQUA principle using isotopically labelled arginine or lysine variants (19), supporting their use as synthetic spike-ins to aid identifications of endogenous peptides. To assess the usability of these peptides, we first determined their linear concentration range, i.e. the range in which changes in quantities generate a proportional change in the recorded MS signal (**Figure 1**).

**Figure 1:**
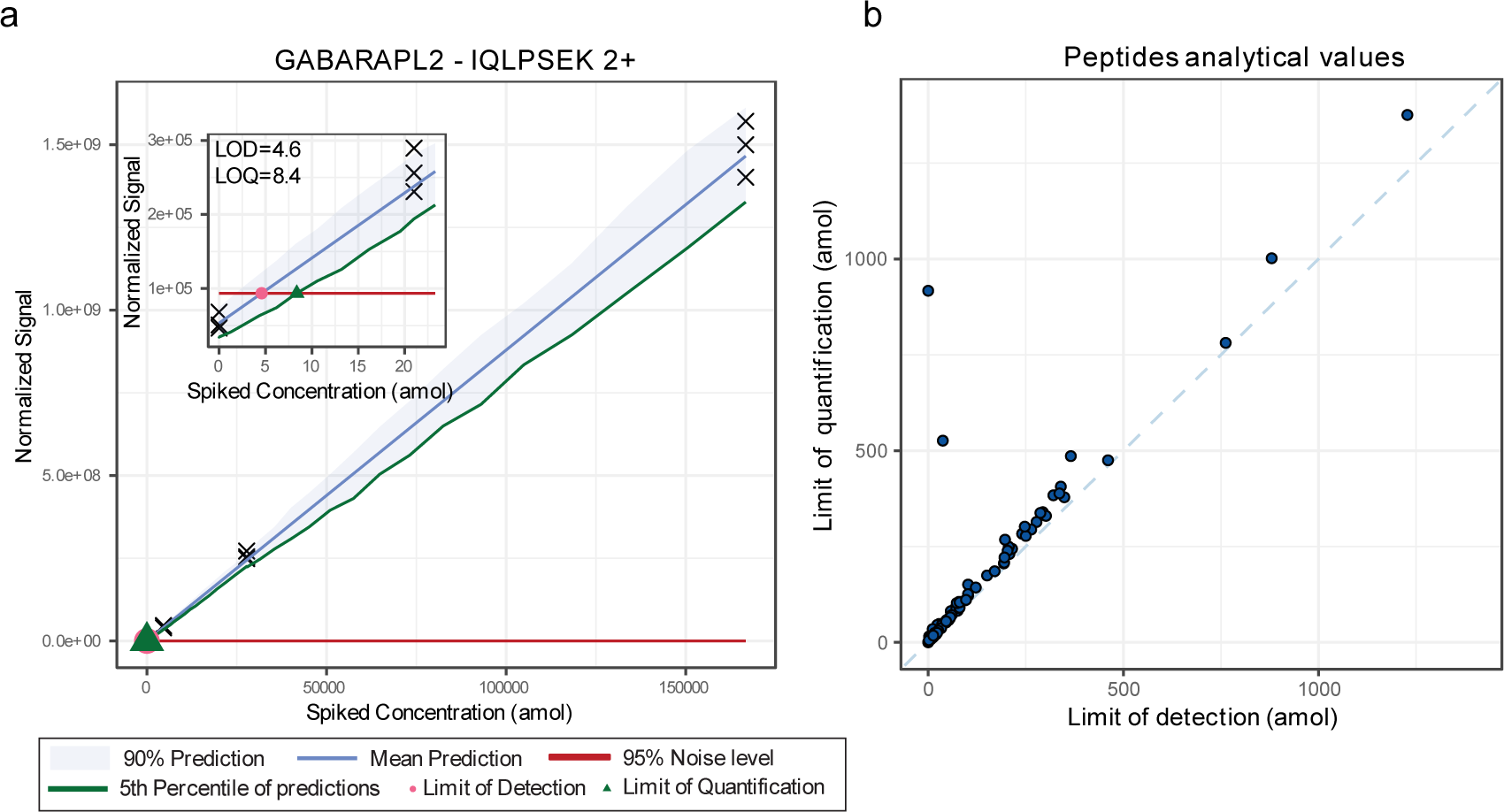
Peptide responses and quantitative accuracy. **(a)** Example calibration curve of the GABARAPL2 peptide IQLPSEK which was detected as doubly charged precursor (2+) and used to monitor GABARAPL2 protein abundance. Inlay is a zoom of the low concentration range. Black crosses indicate measured data points. **(b)** Distribution of limits of detection and quantification of the 94 tryptic peptides used to monitor autophagy.

As with all analytical techniques, quantitative accuracy can only be achieved within a certain analyte concentration range. Quantitative accuracy of targeted proteomics assays is achieved by determining the linear range of detection, the limit of detection and the limit of quantification of respective analytes, *i.e.* the analyte concentrations at which the method generates a signal clearly separated from the noise and a reproducible signal, respectively. These values are peptide-specific and depend in part on the way peptides ionize; hence, they must be determined experimentally for all the target analytes by measuring calibration curves using concentration gradients. For this we followed the envisioned workflow of the assay with the exception that synthetic peptides were spiked after protein digestion (see Methods for details). Briefly, A549 human lung carcinoma cells were harvested, whole cell lysate was generated, and proteins digested overnight using trypsin. To different aliquots of the same starting material, *i.e.* 30 μg of trypsin-digested proteins of whole cell lysate, different amounts of synthetic peptides was spiked, and samples were analyzed by quantitative LC-MS/MS (see Methods for details

The calibration curves for the targeted peptides were measured in three replicates across 7 different concentrations from 6-fold serial dilutions (0.021 fmol to 1’000 fmol on column) and three blank samples. The peptides amounts mentioned throughout this manuscript were calculated based on product weight used for preparing the solutions. Peptide purity was however not 100% and therefore actual values are likely to be lower. As we do not aim for absolute quantification and as these values are only used comparatively between measurements, we provide quantities as indicative values. To only consider the technical variations introduced by the LC-MS/MS setup and not the sample preparation protocol, the triplicate measurements were performed using the same samples. Each replicate of the calibration curve was run in increasing concentration order and blank samples were ran before starting the next curve to limit carry over effects. Only peptide precursors for which a minimum of three fragment ions were detected were used for further analyses. Thus, per peptide a minimum of three LC-MS/MS elution profiles were used for quantification. For most measured peptides, the highest measured concentration (1 pmol on column), caused a loss of signal linearity and/or produced tails in the chromatograms. For this reason, this concentration was not considered for the fitting of the calibration curve. The remaining points were used to fit a calibration curve using the MSStats LOB/LOD package (20). Briefly, this method enables to fit a nonlinear regression model to the intensity-concentration response curve. Limits of quantification (LOQ) and detection (LOD) can then be derived from this model. LODs were defined as the concentrations corresponding to the intersection of the upper bound of the intensity prediction interval for the blank samples (corresponding to the noise level) and the mean predicted intensity of spiked amounts. LOQs were set as the concentrations corresponding to the intersection between the upper bound of the predicted noise intensity and the lower bound of the confidence interval of predicted intensities for given concentrations (see **Figure 1** as example). Out of the 114 considered peptides, 20 could either not be reproducibly detected or accurately quantified and were removed from further considerations, leading to a final list of 94 peptides from 37 POIs with quantified LODs and LOQs (**Figure 2, supplemental Table S2**).

**Figure 2:**
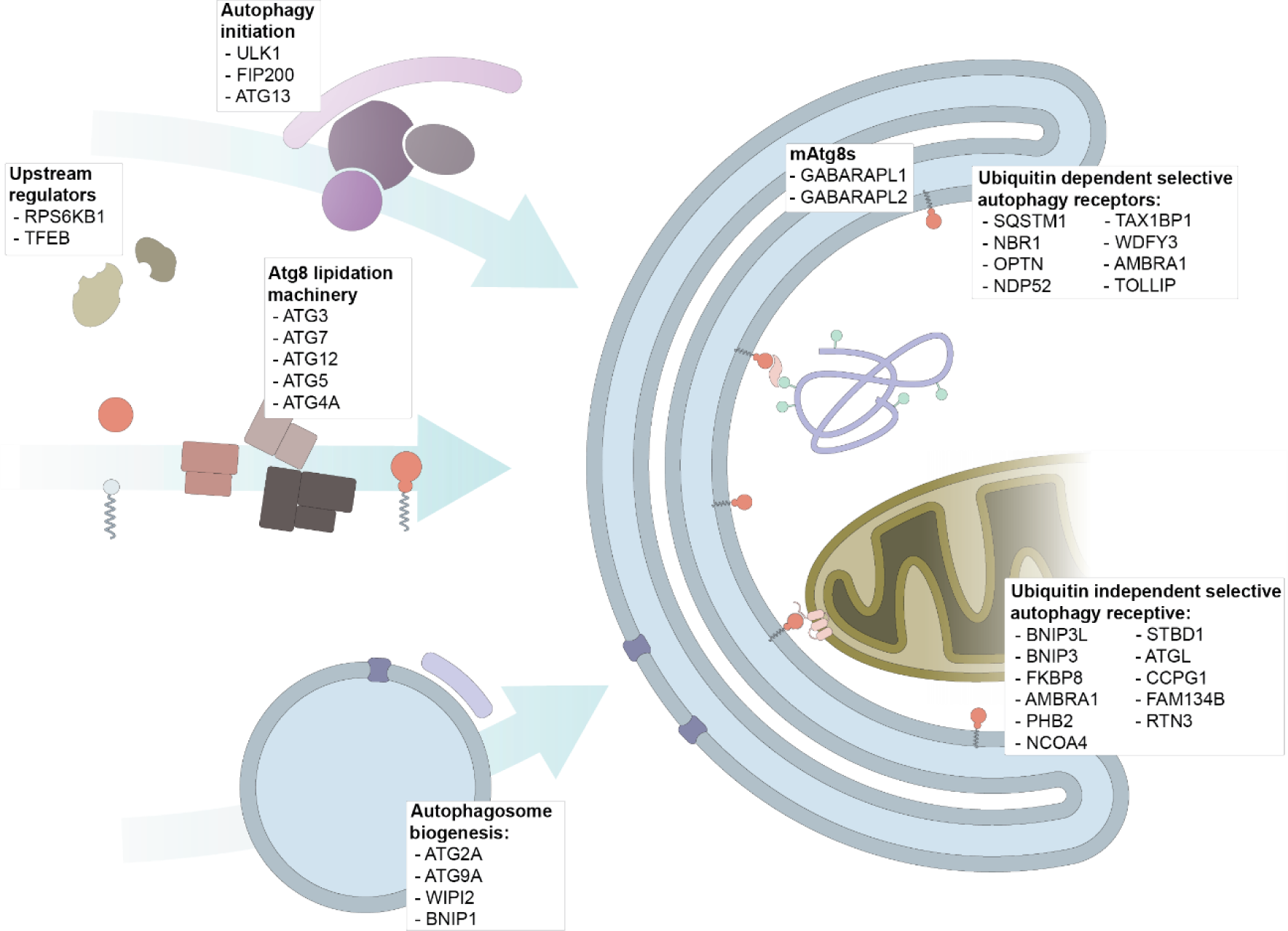
Selected protein targets. Schematic representation of autophagy initiation, highlighting the selected target proteins and their function in autophagy.

The chosen targets include proteins relevant to autophagy regulation (RPS6KB1 and TFEB), the initiation complex (ULK1, RB1CC1/FIP200, ATG13), autophagosome biogenesis (ATG2A, ATG9A, WIPI2, BNIP1, WBP2), the ATG8 lipidation machinery (ATG3, ATG7, ATG12, ATG5, ATG4A) and mammalian ATG8s (GABARAPL1, GABARAPL2), as well as SARs (SQSTM1, NBR1, OPTN, NDP52, TAX1BP1, Alfy/WDFY3, CCPG1, FAM234B, RTN3, NIX/BNIP3L, TOLLIP, FKBP8, BNIP3, AMBRA1, PHB2, NCOA4, STBD1, ATGL, NUFIP1, TNIP1) (**Figure 2**).

### Robustness of the assay compared to standard MS workflows and in a biological context

Amino acid starvation is a strong and commonly used autophagy inducing stimulus. Coupling it to Bafilomycin A1 (BafA1) treatment, which inhibits V-type proton-ATPases blocking lysosomal degradation (5), enables to measure the accumulation of autophagosomal cargo that would be degraded in the absence of BafA1 treatment. To assess the reproducibility of our approach and compare it to standard shotgun proteomics approaches, A549 cells were starved for amino acids in Hank’s Balanced Salt Solution (HBSS) with and without BafA1 for 2 h, samples processed as outlined above, and resulting peptide mixtures were measured on the same LC-MS/MS system comparing PRM-based targeted proteomics to Data-Dependent Acquisition (DDA)-, and Data-Independent Acquisition (DIA)-based discovery proteomics (n=3 technical replicates per treatment).

Out of the 37 protein targets of the assay, 31 could be identified and quantified with a signal above the LOQ in a minimum of one sample using PRM. Standard DIA measurements identified 17 of these proteins and DDA identified 9 (**Figure 3a**). When it comes to quantification, DDA measurements resulted in large coefficients of variation (CVs). Normalized DDA measurements using the MaxLFQ algorithm (21) performed surprisingly well with an average CV of 8.5%; however, only 7 proteins were retained and depending on the protein the CV might be significantly higher (max. value of 44%). Not surprisingly, targeted proteomics developed with specific targets in mind outperformed the other approaches when measuring those targets. In terms of reproducibility, PRM measurements produced an average CV across conditions of 4.8% with most proteins being below 10% (**Figure 3b**).

**Figure 3:**
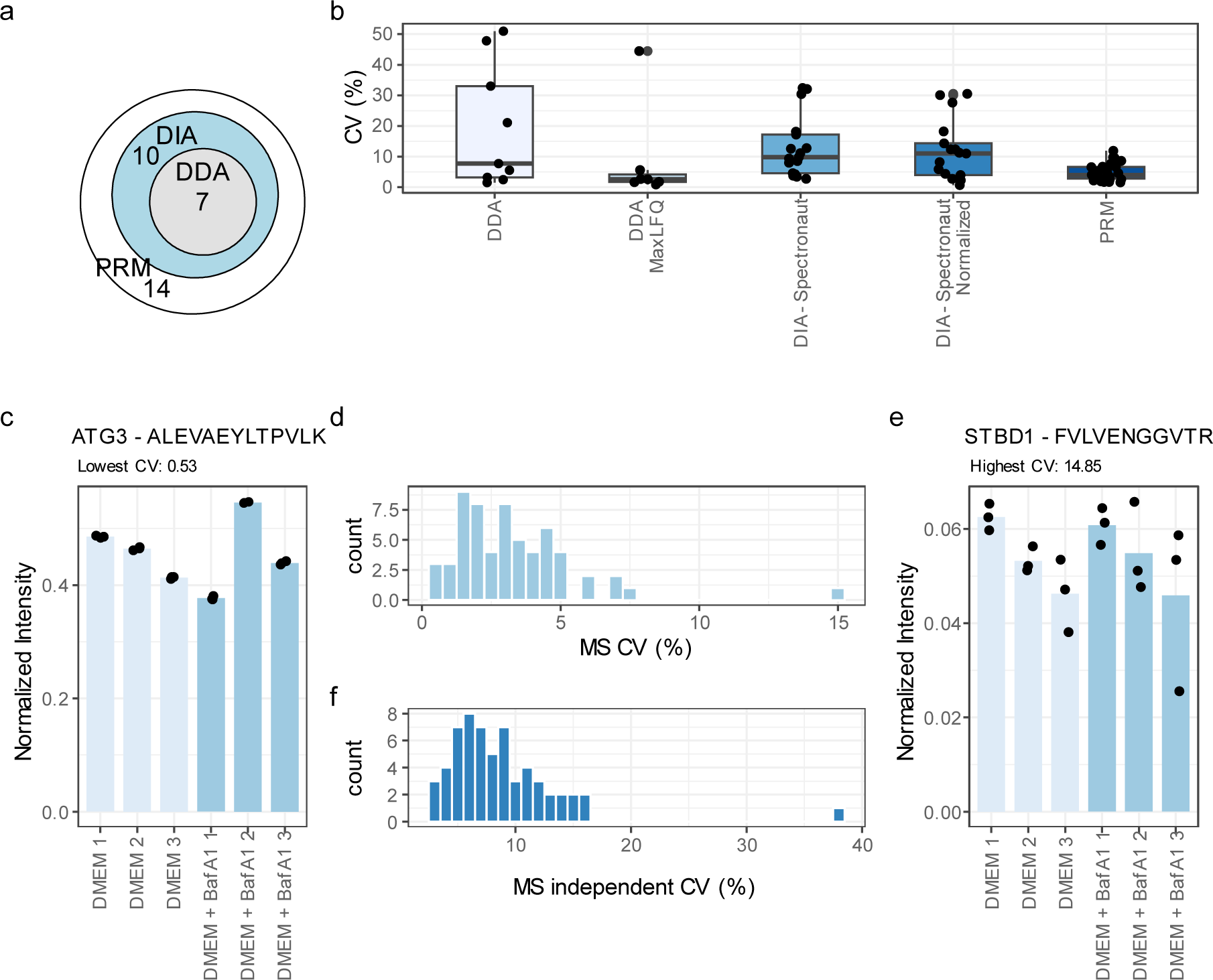
Methods comparisons and measurements of CVs. **(a)** Euler plot of target proteins quantified in a minimum of one condition in DIA (Spectronaut normalization), DDA (MaxLFQ) and PRM. **(b)** Average coefficients of variations at the protein level across HBSS and HBSS + BafA1 treatments using different quantification methods. Single values are represented by black dots. **(c-e)** Average peptides coefficients of variations distribution within biological replicates (d). Detailed data from both lowest and highest CV producing peptides derived from (c) ATG3 and (e) STBD1, respectively, are shown as examples. **(f)** Peptide CVs across biological replicates, representing biological variation and variation introduced by sample preparation.

We then tested the reproducibility of the PRM assay on a larger sample number, using cells harvested in growth conditions (DMEM) in the absence and presence of BafA1 for 2 h. Biological triplicates were prepared for these conditions and all of them were measured 3 times to clearly separate the noise originating from biological differences and sample preparation from the purely technical MS-related noise. For these quantifications, 60 peptides which were identified without interferences in all samples and which gave rise to a signal above the LOQ were considered for CV calculations. CVs calculated for all technical triplicates and averaged across samples ranged from 0.5% to 14.8% with a majority below 5% showing high technical reproducibility of the assay with variations lower or in the range of the variation observed across biological replicates (**Figure 3c-f**).

### Assessment of stimulus-dependent protein degradation by autophagy

Having established the technical principles for our assay, we next asked if we could identify differences in autophagy-dependent protein regulation comparing 2 h amino acid starvation and 2 h glucose starvation to growth conditions in complete media (DMEM). These treatments were coupled to BafA1 treatments for 2 h to study the autophagy flux in all three conditions.

Some conditions like amino-acid starvation (HBSS) considerably reduced protein levels of autophagy cargoes making it difficult to detect and/or quantify these peptides. To calculate protein quantities, we used the following strategy: if a set of peptides was identified in all samples and resulted in a signal above the LOQ, only this set was used for inferring the protein amount. If no valid peptide was found across the whole experiment the protein was considered missing and not quantified. Lastly, if a protein was identified with a given valid peptide set in some samples but none of these peptides were valid in other samples, the protein abundance was imputed with the same standard deviation as the one observed for measured samples and a downshifted mean of 20%. We regard this as a rather conservative estimation of protein abundance in the case of missing values and it can easily be adjusted based on the biological setting of the experiment. This strategy led to the quantification of 34 proteins with peptides above the LOQ.

To identify potential patterns of differential regulation of these proteins across the tested conditions, we filtered the results for proteins showing a significant difference in abundance between treatments (p-value < 0.01 in one-way ANOVA). This filtering yielded a total of 24 proteins. Upon clustering of the results, experimental conditions could be clearly separated, and three major clusters emerged (**Figure 4a**). Cluster 1 is mostly comprised of components of the core autophagy machinery (**Figure 4a-b**). These proteins do not show degradation patterns in any of the tested conditions but seem to accumulate under amino-acid starvation. The levels of these proteins consistently and significantly increase by about 20% once cells are starved for amino acids, regardless of lysosomal activity (**Figure 4b**, p<0.05, T test).

**Figure 4:**
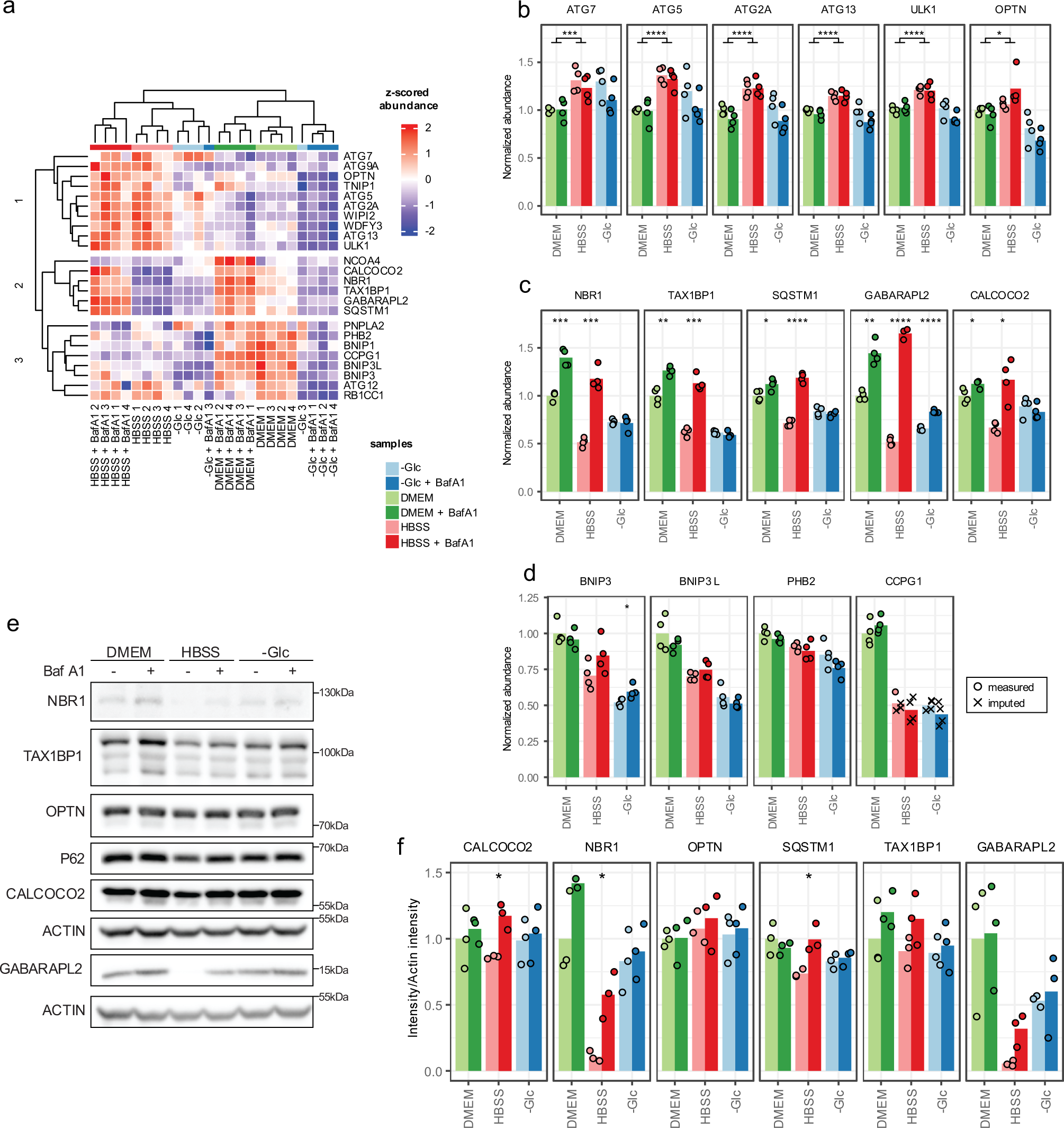
Regulation of protein abundances by autophagy upon different starvation stimuli. **(a)** Heatmap of a hierarchical cluster analysis summarizing the behavior of regulated proteins reaching a p-value < 0.01 in an ANOVA test across all conditions (2 h treatments). **(b-d)** Example proteins for the clusters 1, 2 and 3 from panel a, respectively. Basal conditions (DMEM) were set as 1 for reference. **(e)** Western blot analyses of some of the targets of the proteomics assay. Shown is one representative of n=3 biological replicates. Actin was used for normalization. **(f)** Quantification from the western blots shown in (e). Asterisks represent significance level of a two-side unpaired Student t-test between conditions in (b) and within a condition +/- BafA1 treatment in (c,d, f). *=p<0.05, **=p<0.01, ***=p<0.001, ****=p<0.0001, ns=not significant. Bars highlight average values and dots single experiments.

Cluster 2 constitutes of ubiquitin-dependent, soluble SARs and the ATG8 homolog GABARAPL2 (**Figure 4a, c**). SARs like p62/SQSTM1 are being turned over to some extent under basal conditions and amino-acid starvation drastically increases their flux, corroborating their central role in adaptation to amino acid starvation. In cluster 3 membrane-bound, ubiquitin-independent autophagy receptors BNIP3, NIX/BNIP3L, PHB2 and CCPG1 are not being degraded within lysosomes under any conditions but interestingly seem to be at their most abundant levels under growth conditions (**Figure 4a, d**). The mechanism leading to their decrease under stress conditions is not known. For some proteins, the trends observed by targeted proteomics were also observed by immunoblotting, although the noise levels were higher (**Figure 4e-f**). Interestingly, glucose starvation decreases the overall level of SARs compared to growth conditions. This decrease is independent of lysosomal activity as BafA1 treatment has no effect (**Figure 4c**). The block of turnover is also reflected in the poor clustering of the samples starved for glucose with and without BafA1, as this drug should underscore the autophagy-related cargo degradation. The small but significant stabilization of GABARAPL2 by BafA1 indicates that lysosomes are principally still active, questioning the role of the monitored proteins in autophagy-dependent adaptation to glucose starvation.

### Determination of protein regulation during mitophagy

After demonstrating that our method detects differences in protein regulation within 2 h of amino acid starvation, we wanted to assess whether we could monitor the flux of specific SARs after inducing selective autophagy. We used ARPE-19 cells, a retinal pigment epithelia cell line, treated with the iron chelator DFP to induce selective BNIP3/BNIP3L-dependent and PINK1-Parkin-independent mitophagy (22). These cells expressed the *mito*-QC reporter, a FIS1 mCherry-GFP fusion protein enabling to monitor mitochondrial turnover in the lysosome using fluorescence-based approaches (22). Briefly, this fusion protein emits both red and green fluorescence signals until it is brought to the lysosome, where the acidic conditions quench GFP resulting in a red-only signal that can be detected by flow cytometry or microscopy. We treated cells with DFP for 24 h with and without concanamycin A (ConA), another V-type H+-ATPase inhibitor used to monitor accumulation of autophagosomal cargoes (5). After 24 h of DFP treatment, we observed a significant increase in red mito-lysosomes compared to non-treated control cells by IF microscopy (**Figure 5a-b**). GFP-quenching in lysosomes could be blocked by ConA. Also, flow cytometry analyses confirmed these results (**Figure 5c**).

**Figure 5:**
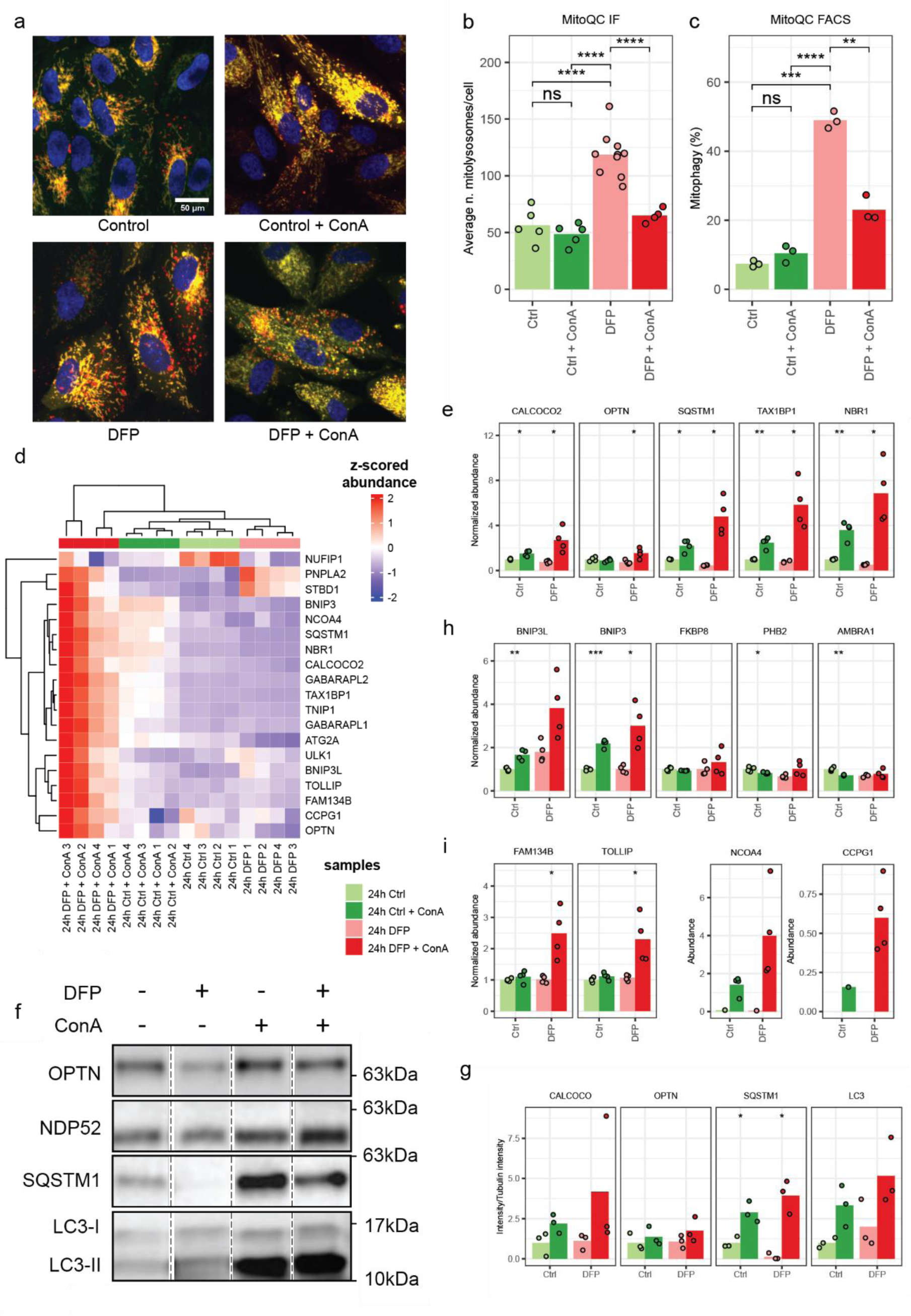
PRM measurements of DFP treated cells. **(a)** Immuno-fluorescence microscopy of *mito*-QC cells. A representative of n=5-10 biological replicates is shown for a total of 200 cells. **(b)** Quantification of the *mito*-QC reporter-based experiments highlighted in (a). Black dots represent the number of mitolysosomes per cell. **(c)** Flow cytometry analysis of *mito*-QC cells. Shown is one representative of n=3 biological replicates **(d)** Heatmap summarizing the changes in protein regulation upon mitophagy induction with DFP. Represented proteins reach a p-value < 0.01 by ANOVA test. **(e)** Detailed quantification of soluble p62-like autophagy receptors linked to mitophagy. **(f)** Western blot analysis of indicated target proteins. Shown is one representative experiment of n=3 biological replicates. Note: dashed lines indicate cropping marks of unrelated samples. All conditions were run on the same gel and blot. **(g)** Quantification of western blots exemplified in (f). **(h)** Detailed quantification of ubiquitin-independent autophagy receptors involved in mitophagy. **(i)** Detailed quantification of other autophagy receptors. Only measured datapoints are represented. If not enough datapoints were measured for DMEM condition (reference point), the samples were not normalized to DMEM. *=p<0.05, **=p<0.01, ***=p<0.001, ****=p<0.0001, ns=not significant, two-sided unpaired Student’s T test. Bars highlight average values and dots single experiments.

Having established that the *mito*-QC reporter cells behave as anticipated, we performed replicate experiments and subjected them to targeted proteomics as outlined above. Protein abundance data were again clustered to gain insights into differential protein regulation. Soluble SARs like p62/SQSTM1, TAX1BP1, NBR1 and CALCOCO2 show a clear autophagy-dependent turnover under growth conditions and increased turnover after DFP treatment (**Figure 5d-e**). Interestingly, OPTN which is linked to PINK1-Parkin-dependent mitophagy, did also slightly respond to DFP treatment (23, 24). These trends were also observed by immunoblotting, although the data was noisier (**Figure 5f-g**).

In contrast to the starvation experiments summarized in **Figure 4**, DFP treatment led also to a turnover of membrane-bound SARs: the mitophagy receptors BNIP3 (25) and BNIP3L/NIX, which is described as a major mitophagy receptor involved in DFP-dependent mitophagy (26), showed low turnover under growth condition but strongly accumulated in DFP treatment and blocked lysosomal degradation (**Figure 5d, h**). AMBRA1, FKBP8 and PHB2 which have also all been linked to mitophagy did not respond to the outlined treatments (27-30). Surprisingly, other receptors also seemed to strongly respond to DFP treatment, like TOLLIP a ubiquitin receptor (31), NCOA4 a ferritinophagy receptor (32), and CCPG1 and FAM134B, two ER-phagy receptors (33, 34). The parallel accumulation of FAM134B and CCPG1 suggests an increased reticulophagy/ER-phagy upon DFP treatment which, to the best of our knowledge has not been described yet, despite DFP being a commonly used iron chelator. This emphasizes the need to comprehensively monitor autophagy receptors to fully grasp cellular responses to stresses.

## Discussion

Autophagy research is a very active field of study and the recent discovery of SARs will undoubtably change our understanding of its physiological relevance. To deepen our understanding of autophagy, these more recent advances need to be considered. Currently, most assays used to study autophagy activity rely on the analyses of single proteins. However, a multitude of selective autophagy subtypes has been discovered (1, 4), which may function in concert. When it is not known which organelle is affected by a given stimulus or pharmacological intervention, it is not trivial to decide which proteins should be analyzed to study the involvement and effects of autophagy. Even if a specific autophagy subtype such as mitophagy is analyzed, the usage of different reporter proteins may yield different results concerning autophagy activity (35). Here, we describe a highly accurate and reproducible, automatable targeted proteomics assay that can be used with low protein amounts in high throughput to screen the autophagic response of cells or tissue specimens in culture. Using this assay, up to 37 autophagy-relevant proteins were identified and quantified, with a focus on SARs.

We would like to point out that the current peptide list is not comprehensive. Since the start of this project, additional SARs such as the golgiphagy receptors YIPF3 and YIPF4, or the lipophagy receptor Spartin have been identified (36, 37), and we plan to include additional peptides to get an even more detailed picture of autophagy regulation. Several hundred peptides can be accurately monitored by current targeted proteomics approaches (12). Importantly, targeted proteomics can also be used to quantify modified peptides, such as ubiquitinated or phosphorylated peptides (38, 39), which will allow to monitor kinase activities and autophagy-relevant signal transduction. In addition, this method does not require genetic manipulations making it suitable for a wide range of samples. While the targets presented here are specific for human samples, the basic principles of this method can be adapted to virtually any organism of interest.

In the current setup, we still rely on the blockage of lysosomal activity to study protein accumulation and to determine autophagy flux. BafA1 or ConA were used for this, which inhibit v-ATPase blocking lysosomal acidification and by this hydrolysis activity (5). To use compounds like BafA1 or ConA, relevant concentration ranges have to be determined which might vary between cell types (40). In yeast, a more robust assay is widely used: the GFP-cleavage assay which monitors the accumulation of protease resistant GFP using GFP-fusion reporters (5). A similar strategy has recently been transferred to mammalian cells monitoring the accumulation of protease-resistant, ligand-bound HaloTag (10). Whereas this assay does not require pharmacological interference with lysosomal activity, it relies on the ectopic expression of tagged marker proteins. We envision that a combination of both approaches, our targeted proteomics approach for screening relevant conditions and a processing assay using the characterized novel targets will yield the most robust assessment of autophagy selectivity and activity. Also, to screen perturbance of autophagy activity in tissue specimens, monitoring of abundance changes of SARs might be sufficient, which would circumvent the need to block lysosomal degradation and would allow analyses of stored specimens, *e.g.* frozen or formalin-fixed, paraffin-embedded tissues, by the outlined approach. More data are needed to robustly address this possibility.

We could demonstrate the suitability of targeted proteomics to measure mammalian ATG8 orthologs, classical markers of so called “bulk” autophagic activity, which are degraded under basal conditions but whose flux drastically increase upon amino acid starvation. Interestingly, it appeared that several proteins of the core autophagy machinery were more abundant under the same conditions. As the observed significant increase is in the range of 20%, such differences would be difficult to detect by expression proteomics or quantitative western blots, highlighting the strength of our approach. As the increase is observable within 2 h of treatment a posttranscriptional regulation likely contributes to the observed abundance changes. Two alternatives come to our minds: (I) ATG proteins could be turned over by a basal non-lysosomal mechanism, which is blocked under stress conditions, *e.g.* the proteasome, or (II) ATG proteins are actively excluded from lysosomal degradation under stress. Since a fixed amount of proteins per assay is used, it could be that “bulk” autophagy is very much upregulated under amino-acid starvation and this leads to a decrease of the overall protein amount whereas the autophagy machinery remains constant, *i.e.* we do not detect an absolute increase in protein amount but a relative increase in the proportion of autophagic machinery in stressed proteomes. Further experiments are needed to answer these questions. Another interesting observation is that glucose starvation blocked all autophagy activity in the used experimental setup. Why this is the case and if this is compensated by proteasomal activity, or is due to reduced transcription/translation is unclear. As lysosomal degradation is commonly regarded as more energy efficient compared to proteasomal degradation (41, 42), a general block in all metabolic activity due to the dependency of cancer cell metabolism on glucose availability seems more likely. Analysis of primary, non-transformed cells may shed more light onto this phenomenon.

We then used the same approach to monitor selective autophagy of mitochondria after iron chelation using DFP. Due to the prolonged timing of this stimulus we observed more prominent autophagy fluxes as in the starvation experiments. Under DFP treatment, ubiquitin-dependent SARs like p62/SQSTM1 and mammalian ATG8 proteins showed clear autophagy activity. In contrast to the starvation experiments, the membrane-bound SARs known to orchestrate this stress response, BNIP3 and NIX/BNIP3L, were identified as clearly upregulated after iron chelation and with a larger flux than under basal conditions. The comprehensive analysis of multiple SARs revealed also a possible increase in ferritinophagy by turnover of the SAR NCOA4. Indeed, DFP treatment was recently shown to also induce ferritinophagy and pexophagy (26, 43). We also identified a potential upregulation of ER-phagy by turnover of CCPG1 and FAM134B upon DFP treatment which, as far as we know, has not yet been described. It appears that DFP treatment leads to a complex cell response, similar to hypoxia having profound influence on the cellular proteome (44, 45). Interestingly, FAM134B was initially reported to be responsible for basal ER sheet turnover (34), which we did not observe here and likely points towards cell-type specific regulation of selective autophagy. These data nicely highlight the need for incorporating multiple receptors into screening experiments. Due to the redundancy of the system, it can be difficult to get a clear readout of organelle fluxes and this approach is best combined with more targeted approaches like the *mito*-QC reporter system, which in our case confirmed the engulfment of mitochondria into lysosomes by DFP treatment.

As most published research focuses on a very narrow subset of receptors, we only have limited understanding about the physiological responses to stress conditions and the interplay of selective autophagic activities. We strongly believe that our approach offers a more detailed picture of the actual response to various stresses within a cell. By automating MS sample processing, the described assay is easily adaptable for high sample throughput (46, 47), supporting e.g. drug screening approaches to characterize pharmacological autophagy modulators. Due to its accuracy, sensitivity, and throughput, we see the applicability of this assays in both basic/preclinical science to study molecular mechanisms of autophagy as well as in clinical science, be it prognosis or diagnosis of health states. The goal will be to identify robust protein biomarkers whose abundance profiles reflect the autophagy state of a given biological specimen.

## Material and methods

### Cell culture

A549 lung carcinoma cells were grown in Dulbecco’s Modified Eagle Medium (DMEM, PAN biotech) supplemented with 10% Fetal Bovine Serum (FBS, BioWest) and Penicilin Streptamycin (PAN Biotech). 2.2×10^6^ cells were seeded into 10 cm dishes to reach about 80% confluency on the next day. Treatments were performed by washing the plates twice with PBS before changing to the treatment medium for 2 h. Treatment media were Hank’s Balanced Salt Solution (HBSS, Thermo Fisher) for amino-acid starvation and DMEM glucose-free (-Glc, PAN biotech) supplemented with 10% dialyzed FBS (BioWest), penicillin/streptomycin (PAN Biotech) and stable glutamine (GlutaMAX, Thermo Fisher). Bafilomycin A1 (Santa Cruz) was used at a final concentration of 2 nM. Cells were washed twice with ice-cold PBS before harvesting by scrapping on ice. Cell pellets were snap frozen in liquid nitrogen.

The ARPE-19 (Human Retinal Pigment Epithelial cell line) stably expressing the *mito*-QC reporter cell lines was generated in the laboratory of Dr. Ian Ganley (22). Cells were grown in Dulbecco’s Modified Eagle Medium low glucose (DMEM, Roth) supplemented with Ham’s Nutrient Mixture F12 (F12, Sigma) 10% Fetal Bovine Serum (FBS, Panbiotech) Glutamine (Roth) and Penicilin Streptamycin (Gibco). 500,000 cells were seeded into P6-well plate to reach about 80% confluency on the next day. Cells were then treated with a mitophagy inductor, DFP (Deferiprone) for 24h and plus-minus a specific inhibitor of vacuolar type H+-ATPase activity ConA (Concanamycin A).

### MS samples preparation

Pellets were lysed in 1% sodium deoxycholate buffer in 50 mM ammonium bicarbonate pH 8.5. Samples were supplemented with 1unit/µl benzonase (Dr. Nuclease Benzonase, Syd Labs). Protein concentration was measured and adjusted using BCA assay and samples were spiked with 12 or 120 fmol of heavy-labeled peptides. Samples were subsequently diluted to reduce sodium deoxycholate below 1%. Samples were reduced by adding 1 mM DTT and incubated at 37°C for 30 min, then alkylated with IAA for 15 min in the dark at room temperature for 15 min.

Samples were digested with trypsin to a 1:100 trypsin (Promega) to protein ratio for 15 h at 37°C with constant agitation. Trypsin was inhibited and sodium deoxycholate was precipitated adding 50% TFA to a final concentration of 2%. Samples were desalted using AssayMAP C18 cartridges (Agilent) on a Bravo liquid handling platform (Agilent), and eluted in 50 µl 80% acetonitrile and 0.1% formic acid. Solvents were lyophilized to remove organic solvents. Peptides were resuspended in 0.1% formic acid to a 1 µg/µl.

### Synthetic peptides

Heavy-labelled synthetic standards for all peptides mentioned in supplemental Table S2 were acquired from SpikeTides (JPT) or custom synthesis (GenScript) with the following chemical modifications: Carbamoylmethylated cysteine, Carboxy-terminal ^13^C_6_-^15^N_2_ Lysines, ^13^C_6_-^15^N_4_ Arginines. Peptides were not purified but isotope label purity was high. Crude peptides were used and the purity of peptides was not taken into account for calculating concentrations.

### LC-MS/MS

LC-MS/MS measurements were performed on an EASY-nLC 1000 nano-flow UHPLC system (Thermo Fisher Scientific) coupled to a Q Exactive HF-X hybrid quadrupole-Orbitrap mass spectrometer (Thermo Fisher Scientific). 5 µl of solubilized peptides in solvent A (0.1% formic acid in water) were separated on a fused silica HPLC column (75 μm internal diameter column Fused-silica PicoTip® emitter: SilicaTip™, New Objective) self-packed with Waters Acquity CSH C18, 1.7 μm (Waters) to a length of 20 cm) using a linear gradient of solvent A and solvent B (0.1% formic acid in 80% acetonitrile in water) from 4% solvent B to 30% over 85 min, followed by an increase to 100% buffer B over 8 min and 7 min at 100% buffer B at a 250 nl/min flow rate. The spray voltage was set to 2.3 kV with a capillary temperature of 250°C.

Mass spectrometer was operated in PRM mode at 60’000 resolution with an AGC target set as 1e6 and a maximum injection time of 118 ms. Isolation window was set at 1.5 m/z, normalized collision energy was set at 27. Data was acquired in centroid mode. Spectral libraries were acquired using a mixture of the synthetic peptides of reference in Data Dependent acquisition with the same resolution and collision energies.

### Data Analysis

Raw data was analyzed using Skyline (48, 49). MS2 spectra were matched to libraries generated using the synthetic peptides. Extracted ion chromatograms were manually inspected, signal from interfering ions were removed and integration peak boundaries were modified if needed. Precursors with less than 3 valid transitions were excluded from further considerations. To normalize the intensity across runs, an additional peptide from the background matrix QSVENDIHGLR from Type I cytoskeletal keratin 18 was monitored for the calibration curve. For experiments, precursor intensities were normalized on their respective heavy-labelled peptides. Fitting of calibration curves was performed using the MSStats LOBD package (20).

Further data analysis was performed using in-house R code. Peptides intensities were calculated as the sum of light fragments divided by the sum of heavy fragments. Protein intensities were calculated as the average intensities of light fragments with heavy counterparts divided by the heavy fragments intensities for all peptides belonging to this protein if they reached the LOQ in a minimum of a replicate. Heatmaps were generated using the Complex Heatmap R Package (50).

### Flow Cytometry

Medium containing floating dead cells was collected in a standard plastic flow cytometry tube. Cells were washed with 500 μL of sterile PBS 1X to remove culture medium and PBS was also collected. 150 μL of 0.05% Trypsin was added per well and cells were incubated for 5’ in an incubator at 37 °C and 5% CO2. 450 μL of FBS-containing complete medium was added to each well to inactivate Trypsin. Cells were collected in the flow cytometry tube and pelleted by centrifugation at 1200 rpm for 5’.

Supernatant was discarded and 100 μL of complete medium without phenol red was added to each tube. Samples were resuspended by gentle shaking and propidium iodide was added to each tube 5 minutes before the flow cytometry analysis, reaching a final concentration of 100 ng/mL, for viable population selection. Tubes were kept in ice until FC analysis. The analysis was performed using a Cytek Aurora 5 lasers (Cytek Biosciences) At least 10,000 events were acquired. Control cells were used to set the threshold of mitophagyhigh population defined by a mCherry/GFP ratio of ∼5%. Analysis was performed using the Spectroflo (Cytek Biosciences) for the umixing of the data and later it was analized in Flowjo v10.10 (BD Biosciences).

### Protein extraction and western Blot

Cells were washed with PBS before harvesting by scrapping with lysis buffer (Tris-HCl 50mM, Glycerol 10%, SDS 2%, protease and phosphatase inhibitors) and boiled for 15’ at 95°C. Protein concentration was determined with the Pierce BCA Protein Assay (23227, Thermofisher) following the manufacturer’s instructions. Protein extract (20 µg) was supplemented with 5X loading buffer (4% glycerol, 0.5 M Tris-HCl pH 6.8, 8% SDS, 0.04% bromophenol blue, 5% β-mercaptoethanol) and resolved on Any kD Criterion TGX Precast Stain-free gels (5678124, Bio-Rad). Proteins were transferred to 0.2 µm PVDF membranes using a TransBlot Turbo Transfer System (Bio-Rad). Membranes were blocked with 5% non-fat milk in PBS-T (0.5% Tween-20 [1706531, Bio-Rad] in PBS) for 1 hour. Membranes were then incubated overnight at 4°C in primary antibodies diluted 1:1000 (Table S1) in 3% dry milk in PBS-T + 0.02% sodium azide, and subsequently for 1 hour at room temperature in secondary antibodies diluted 1:5000 in 3% dry milk in PBS-T. Membranes were developed using Pierce ECL Western Blotting substrate (32106, Thermo Fisher) and in ECL select Western blotting (GERPN2235, Cytiva). Following antibodies were used: anti-NDP52 (Atlas, HPA023295), anti-LC3 (Nanotools, 0231-100/LC3-5F10; Novus, NB100-2220), anti-TAX1BP1 (Sigma, HPA02432), anti-SQSTM1 (Cell Signaling, 5114S; Abcam, ab56416), anti-NBR1 (Cell Signaling, 5202S; Santa Cruz, SC-130380), anti-OPTN (Santa Cruz, SC-166576), anti-beta Actin (Santa Cruz, SC-47778HRP).

### Assessment of mitophagy by fluorescence imaging

Spinning disk confocal microscopy (VisiScope CSU-W1, Visitron System GmbH) was used for *in vivo* imaging. Nuclei were stained with Hoechst and the images were acquired with Photometrics pco.edge 4.2 sCMOS camera, a 40x objective and 0.35 µm z-step. Images with the *mito*-QC reporter to quantify mitophagy were acquired using a Leica SP5 laser scanning confocal microscope (HC PL APO 63x/1.40 oil CS2). All the images were processed with FIJI v1.54f software (ImageJ, NIH). Quantification of mitophagy was performed from five to ten independent fields counting over 200 cells for condition. Images were processed with the *mito*-QC Counter, as described in (51). For images acquired the following parameters were used: Radius for smoothing images=1, Ratio threshold=0.5, and Red channel threshold=mean + 0.5 standard deviation.

## Data Availability

MS raw data and skyline files are available from Panorama (https://panoramaweb.org). R scripts used for data analysis and figure generation are available from GitHub (https://github.com/DengjelLab/Autophagy-Readout-PRM). Fragment ions, western-blots, immunofluorescence and FACS quantifications used as input data for figure generation and statistical testing are available from the same repository.

## Abbreviations

ATG: autophagy related
CV: coefficients of variations
ER: endoplasmic reticulum
LC: liquid chromatography
LOD: limit of detection
LOQ: limit of quantification
MS: mass spectrometry
PRM: parallel reaction monitoring
SARs: selective autophagy receptors

## Acknowledgements

The authors wish to thank Sarah Cattin from the Cell Analytics Facility for assistance with FACS analysis, Felix Meyenhofer from the Bioimage Core facility for imaging support, and Anne Oberson for technical assistance. This work was supported by the Canton and the University of Fribourg as part of the SKINTEGRITY.CH collaborative research project and by the Swiss National Science Foundation (grant CRSII5_189952 to JD and 310030_215271 to PB).

## Supplemental Material

**Supplemental Table S1:**
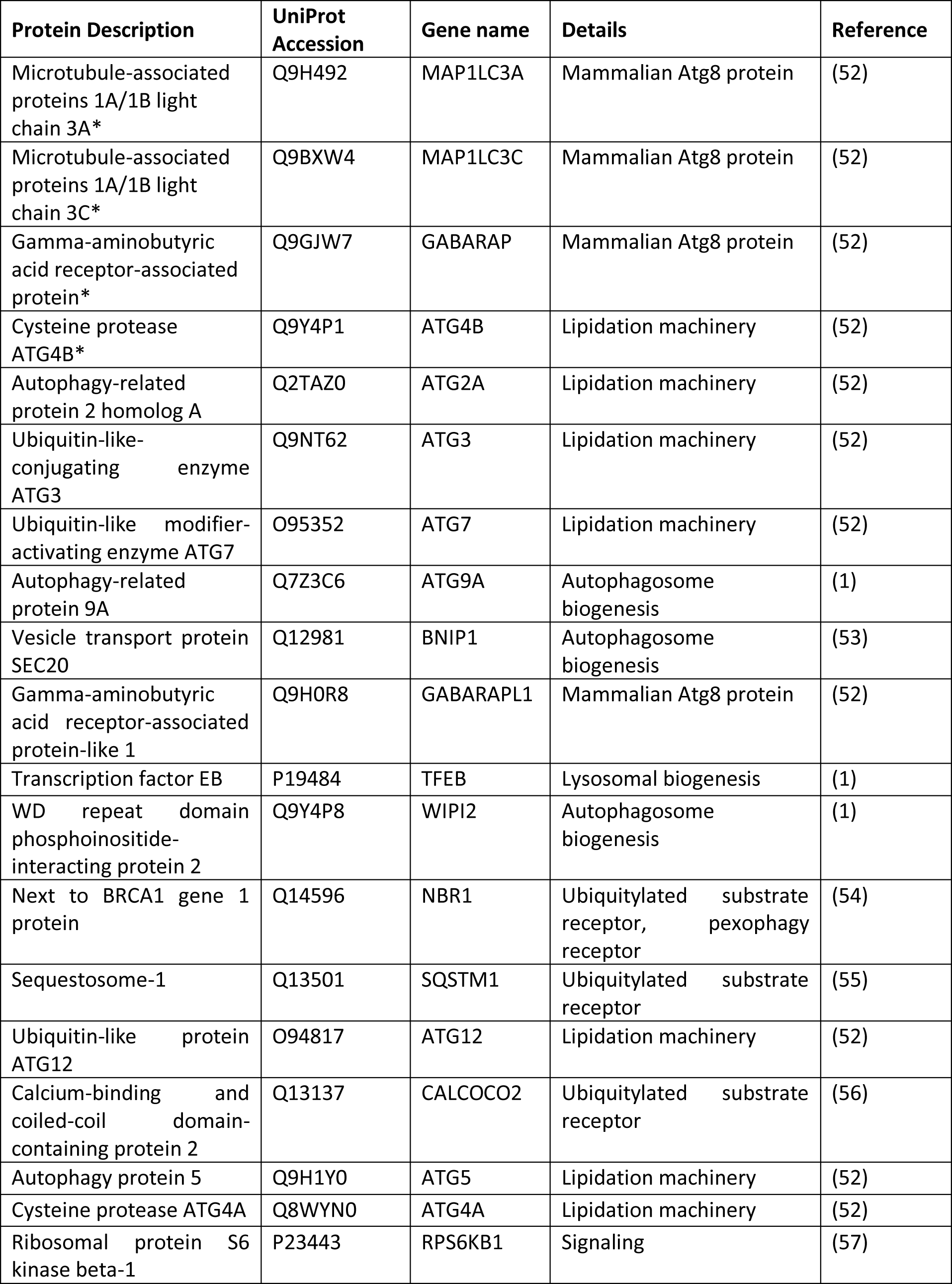
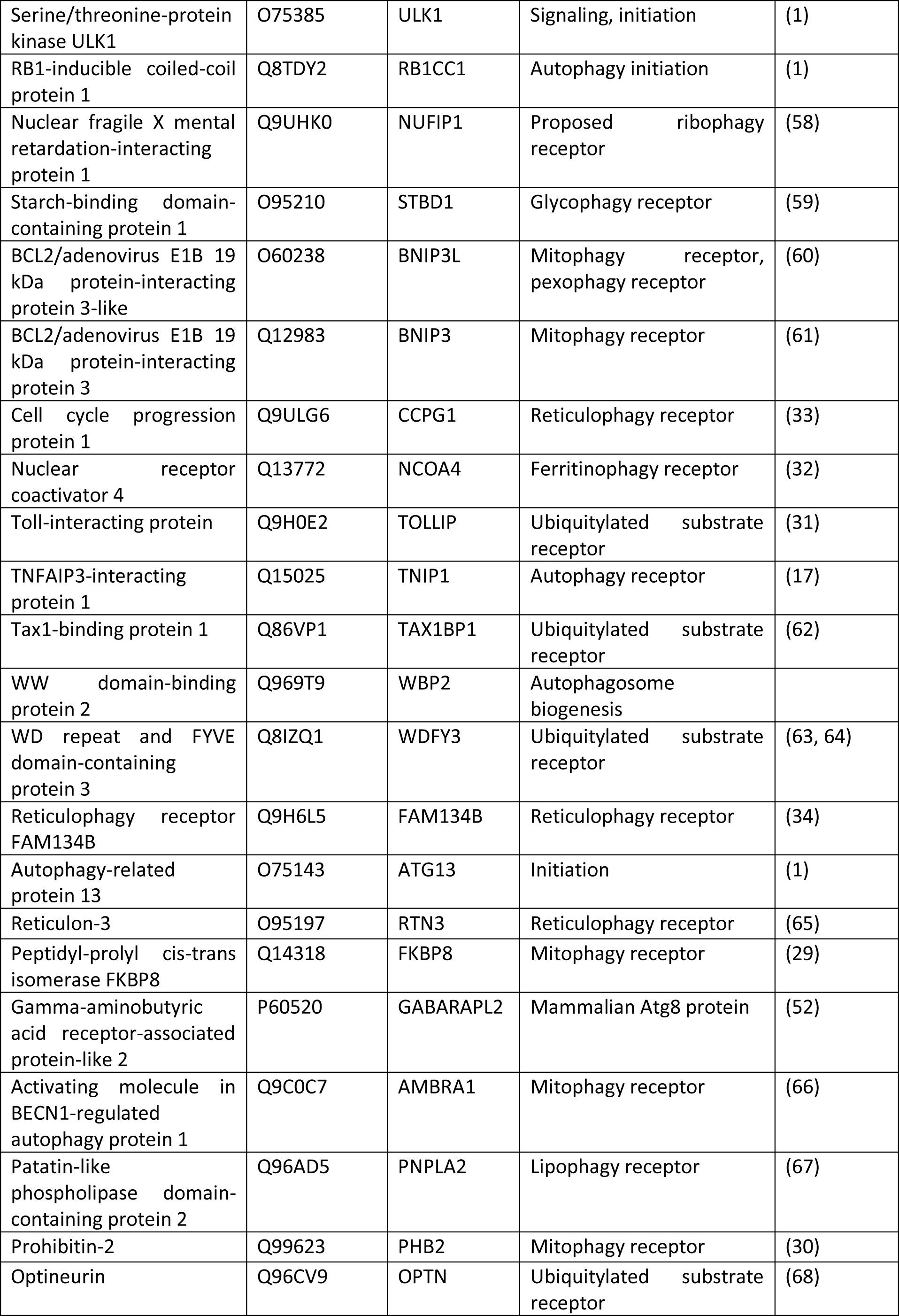
Target proteins monitored by PRM.

**Supplemental Table S2:**
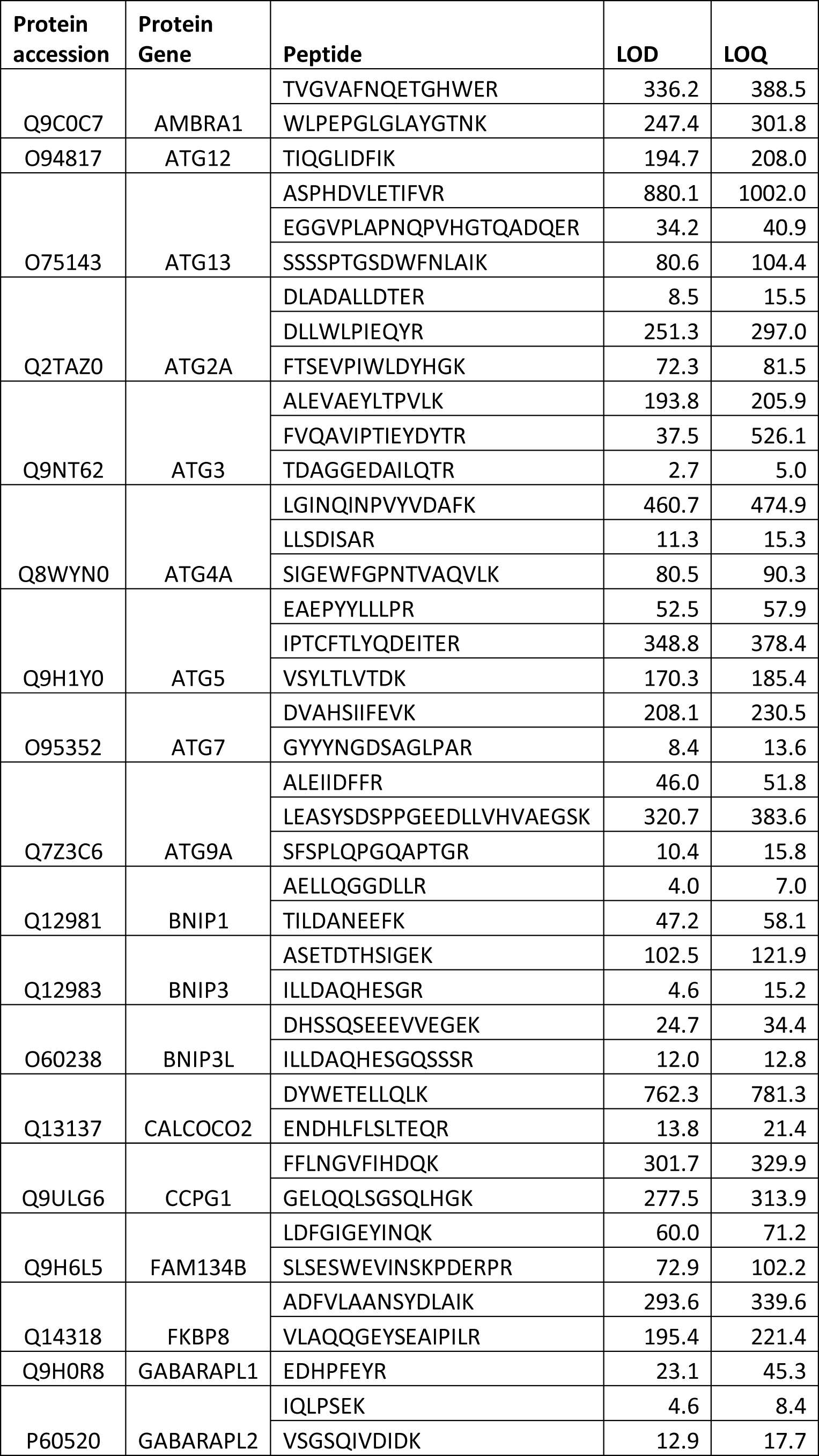
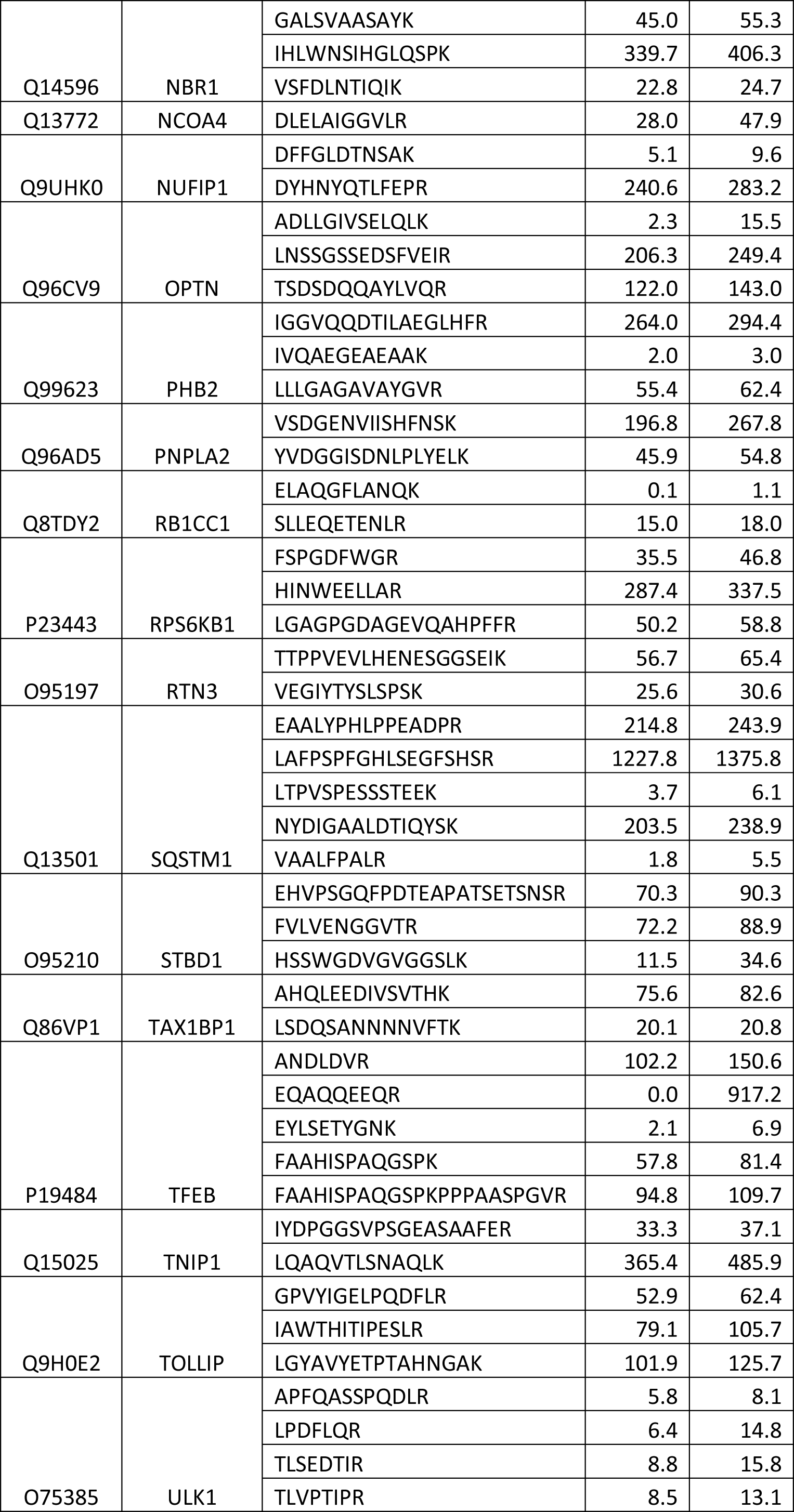
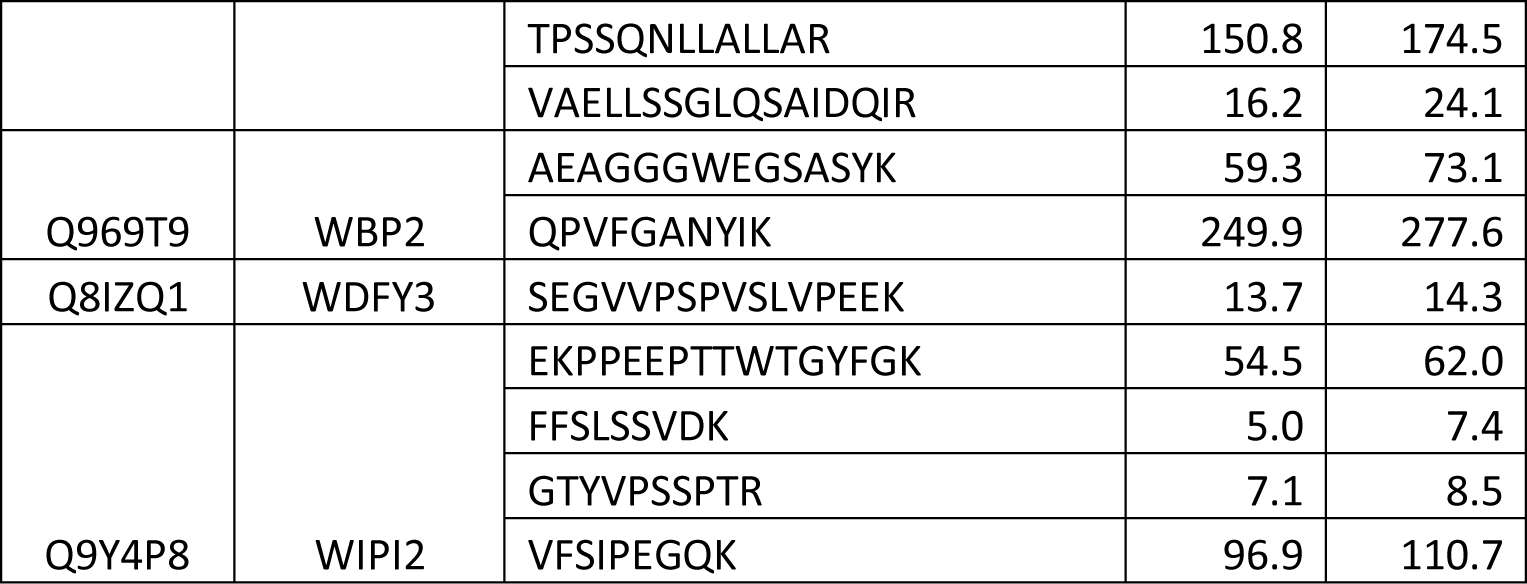
Validated peptides and their analytical figures of merit.

